# Direct Genome Transfer from *Acholeplasma laidlawii* to Yeast

**DOI:** 10.1101/2025.10.17.683135

**Authors:** Daniel P. Nucifora, Tanveer S. Saini, Bogumil J. Karas

## Abstract

Cloning bacterial genomes in yeast or other hosts is a crucial step in workflows for creating highly engineered strains. The genome of *Acholeplasma laidlawii* can be cloned in yeast if a toxic gene is removed, but this deletion has not been tested in live bacteria. Here, we demonstrate cloning the 1.46 Mb genome of *A. laidlawii* strain DN-E by integrating a yeast vector into the toxic gene and transferring the genome directly to yeast using cell fusion. Genomic integration was only possible when *A. laidlawii* was complemented with a functional recA gene. Following cell fusion, 13 out of 20 screened yeast colonies contained the *A. laidlawii* genome, as determined by multiplex PCR. This is the first demonstration showing cloning entire *A. laidlawii* genomes by simply mixing donor and recipient cells in fusion buffer, bringing us one step closer to creating fully synthetic *Acholeplasma* strains.

**GRAPHICAL ABSTRACT:** 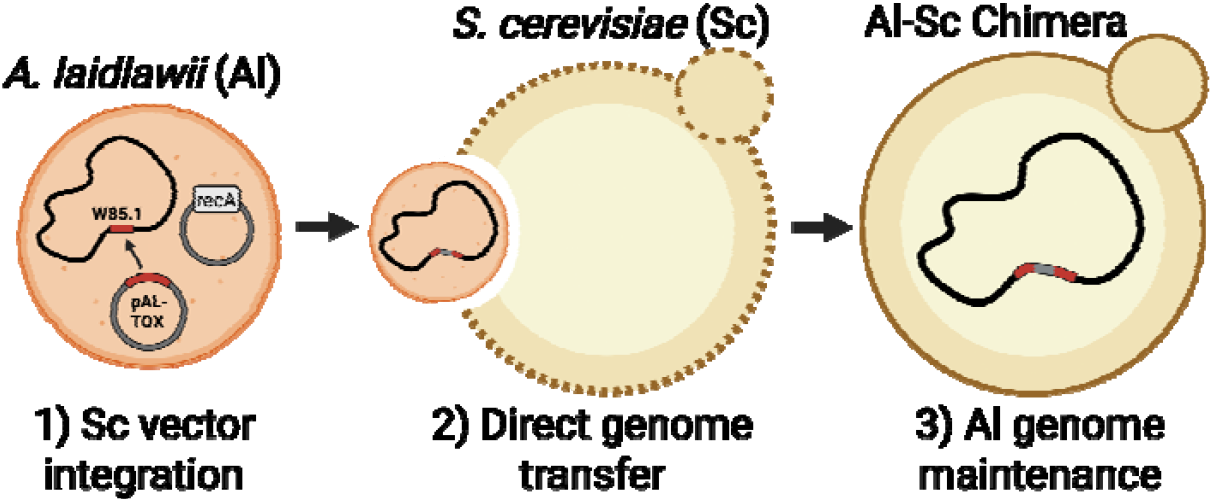

## INTRODUCTION

The baker yeast *Saccharomyces cerevisia*e is widely used as a platform for synthetic genomics due to its highly efficient homologous recombination machinery and its ability to stably propagate very large DNA constructs, including those exceeding one megabase. These features have enabled the capture of complete bacterial genomes in yeast, from the ∼0.5 Mb minimal *Mycoplasma mycoides* JCVI-syn3.0 genome to nearly 2 Mb genomes such as *Haemophilus influenzae* and *Spiroplasma citri*^1–3^.

In several cases, an approach known as genome transplantation has been used to reintroduce yeast-maintained genomes into recipient bacteria, effectively reactivating them as living cells^4,5^. Modification of cloned genomes in yeast can result in transplanted bacteria with new genotypes and phenotypes^5^. When combined with the extensive genome editing tools available in yeast^6–9^, this strategy establishes a powerful framework for redesigning and engineering bacterial strains at the whole-genome level.

Cloning whole genomes in yeast requires the addition of elements that constitute a yeast centromeric plasmid (YCp): a centromere, an autonomously replicating sequence (ARS), and a selectable marker^10^. Low-G/C prokaryotic genomes are more likely to contain endogenous sequences that can act as ARSs in yeast, allowing for their replication without additional ARS installation that would otherwise be necessary in higher-G/C sequences^11^. As described previously, three main strategies have been established for cloning bacterial genomes in *S. cerevisiae*^10^. In transformation-associated recombination (TAR) cloning, an isolated genome is co-transformed into yeast with a YCp vector that is designed to recombine into the genome^10^. In a related method, the whole genome is instead assembled from recombination between overlapping fragments^10,12^. These methods do not require any bacterial genetic tools, but if tools are available, the vector can instead be inserted in live bacteria, followed by whole-genome isolation and transfer to yeast^10,13^.

It has since been shown that live bacteria harboring a genomic YCp vector can also be directly transformed into yeast spheroplasts using polyethylene glycol^3^. This method, termed cell fusion, enables transfer of genomes from bacterial to yeast without the need for any DNA isolation, simplifying the cloning approach and protecting genomes from shearing^3^. Cell fusion has also been coupled with CRISPR-Cas9 to enable both cloning and editing of bacterial genomes within *S. cerevisiae* ^*14*^. Despite the name of this method implying membrane fusion, it is known that at least some bacteria are engulfed by yeast through endocytosis^15^.

Many genomes that have been cloned in yeast are from species in the *Mollicutes* class, which comprises cell-wall-lacking bacteria with small (0.58 – 2.2 Mb^16^) A/T-rich genomes. Genome transplantation is currently limited to only a few *Mollicute* species, specifically to those within the Spiroplasma phylogenetic group^1,17^. We have previously expressed our goal to create synthetic strains of the *Mollicute* species *Acholeplasma laidlawii*, for which transplantation of yeast-cloned genomes is a crucial step. The 1.5-Mb genome of *A. laidlawii* strain PG-8A has previously been cloned in yeast using TAR cloning^18^. One gene in the *A. laidlawii* genome, ACL_0117, was found to be toxic to yeast, and cloning was only successful when the YCp vector recombined into the genome to replace this gene^18^. Although the issue of toxicity had been overcome, it is currently unknown whether this cloned *Acholeplasma* genome is viable; integration of the vector and deletion of ACL_0117 occurred within yeast and were not tested in bacteria. Furthermore, the reported cloning approach was inefficient, labor-intensive, and required experienced researchers to isolate and transform the entire *A. laidlawii* genome to yeast.

These limitations have motivated us to replicate cloning of the *A. laidlawii* genome in yeast by instead using cell fusion with a strain carrying an integrated YCp vector. This approach will ensure viability of the genome before it is cloned in yeast. Furthermore, cell fusion should be more efficient compared to TAR cloning, since successful cloning is not contingent on co-delivery or recombination of the genome and vector in yeast. However, genomic integration of a vector is not practical for strain PG-8A, given our observations that it is highly resistant to transformation. We therefore chose to use the related strain *A. laidlawii* 8195 for this work, as we have developed genetic tools for this strain, including a new electroporation protocol, new shuttle vectors, and a protocol for delivery of Tn5 transposomes^19^. We specifically used our evolved 8195 strain, *A. laidlawii* DN-E, as it is faster growing and more receptive to plasmid transformation.

The DN-E genome contains a gene encoding for Protein Accession ID WIF88585.1 that shares 95% nucleotide identity with ACL_0117 from strain PG-8A. Given the reported toxicity of ACL_0117^18^, we expect that the DN-E homolog (hereby referred to as W85.1) will also need to be inactivated prior to cloning in yeast. A targeted approach using homologous recombination could be used to transform *A. laidlawii* and integrate a YCp vector directly into the toxic gene to disrupt it, assuming it is not essential. Targeted integration of a non-replicative plasmid has been demonstrated for *A. laidlawii* strain JA1, which is an ancestral strain of 8195 and DN-E^20,21^. However, it is known that strain 8195, and subsequently strain DN-E, contain a nonsense mutation in the recA gene that results in a truncated protein product; this mutation is thought to abolish DNA repair and recombination^20,22^. Thus, as strain DN-E exists currently, targeted integration using homologous recombination would likely be unsuccessful.

We thought that it should be possible to restore recombination in strain DN-E through complementation with a functional recA gene. The inclusion of the *Escherichia coli* recA gene in non-replicative plasmids has been shown to improve rates of genomic integration in *Mycoplasma* species^23,24^. We instead chose to use recA from *A. laidlawii* strain PG-8A for complementation, as the PG-8A recA gene is nearly identical to that of strain JA1 on the amino-acid level, and it does not contain the nonsense mutation found in strains 8195 and DN-E. Here, we demonstrate that recA complementation in *A. laidlawii* strain DN-E enables targeted integration of a non-replicative plasmid into the putative toxic gene W85.1. To facilitate this integration, we developed additional genetic tools, including lacZ expression using an endogenous promoter from strain PG-8A, and we assessed plasmid stability using the shuttle vector pNZ18-CAH. Finally, we show that the modified *A. laidlawii* genome, carrying an integrated yeast plasmid, can be directly transferred into *S. cerevisiae*, enabling whole-genome cloning in yeast.

## RESULTS & DISCUSSION

### Expression of lacZ in A. laidlawii

Currently, TAR cloning is the only method that has been established for cloning the *A. laidlawii* genome in yeast^18^ (Figure 1A). To establish vector insertion for cell fusion, we needed to sequentially transform *A. laidlawii*, first with a recA-complementation plasmid, and then an integrative plasmid targeting the putative toxic gene W85.1 (Figure 1B). We aimed to clone the PG-8A recA gene into one of our shuttle vectors, pAL1 or pNZ18-CAH^19^, for delivery to strain DN-E. However, it was first important to verify that these vectors can be used for exogenous gene expression in *A. laidlawii*. We opted to test this using the reporter gene lacZ, as expression is easy to verify with a blue-white screen, and lacZ is known to function in the related species *Acholeplasma oculi*^25^. Our initial attempts to express lacZ in strain DN-E using *E. coli* Anderson promoters were unsuccessful, leading us to first identify a strong constitutive promoter for *A. laidlawii*.

**Figure 1.**
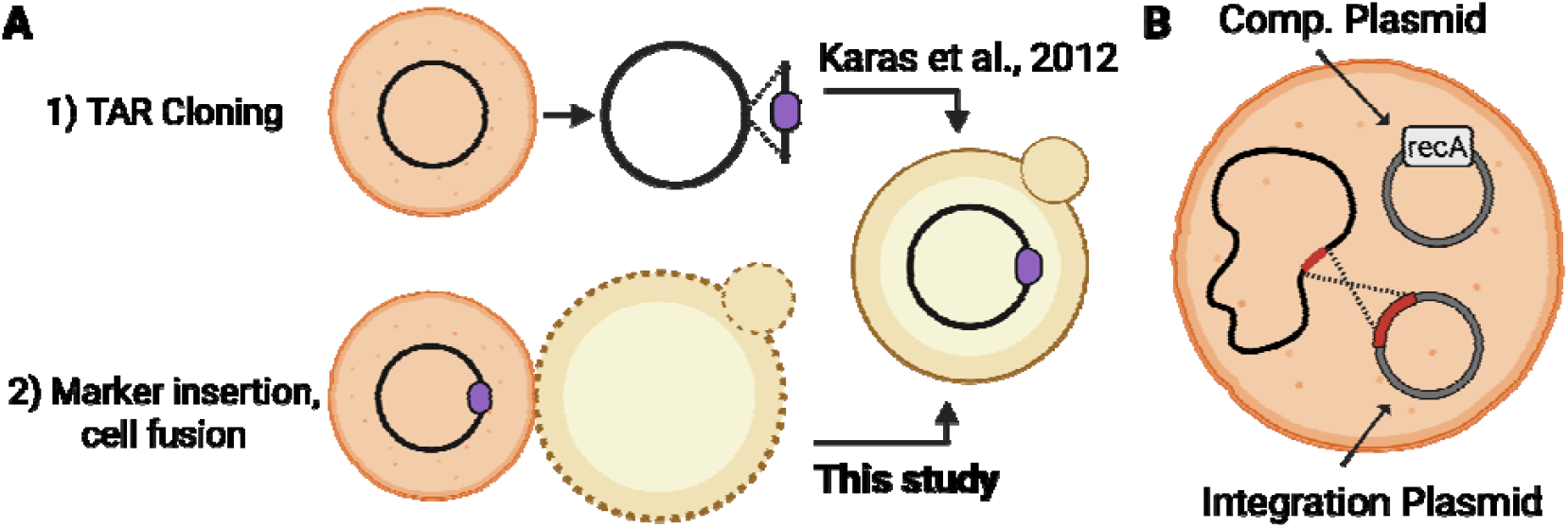
Strategies for cloning the *A. laidlawii* genome in yeast. (A) Two strategies for cloning whole genomes derived from *A. laidlawii* to *S. cerevisiae*. Strategy 1 has been demonstrated, Strategy 2 was proposed for research presented in this manuscript. Yeast elements are represented as a purple oval^10^. (B) Proposed strategy for integration of yeast elements in *A. laidlawii* DN-E through two subsequent transformations. First, a complementation plasmid will be introduced to restore homologous recombination. Next, a non-replicative plasmid will be transformed to integrate directly into the putative toxic gene W85.1.

Endogenous *A. laidlawii* 8195 promoters have previously been identified by cloning randomly digested genomic fragments in *E. coli* to drive expression of alpha amylase^26^. We instead referred to transcription-start-site (TSS) data available for strain PG-8A^27^, where we selected the rpsJ operon as a relatively highly expressed transcript. We PCR-amplified a 368-base-pair fragment upstream of the rpsJ TSS from the PG-8A genome to capture this promoter. This fragment was cloned into pAL1 and pNZ18-CAH, upstream of a promoter-less lacZ gene containing an *E. coli* terminator, to create pAL1-Z and pNZ18-Z, respectively (Figure 2A). *E. coli* colonies carrying pAL1-Z and pNZ18-Z were blue, and transformation of pAL1-Z to *A. laidlawii* DN-E yielded blue colonies when SP-4 plates were supplemented with X-gal (Figure 2B). However, when we transformed pNZ18-Z to *A. laidlawii*, we only observed white colonies on X-gal-supplemented plates. Recovery of pNZ18-Z from *A. laidlawii* back to *E. coli* also yielded only white colonies, suggesting inactivation of the lacZ gene on our plasmid.

**Figure 2.**
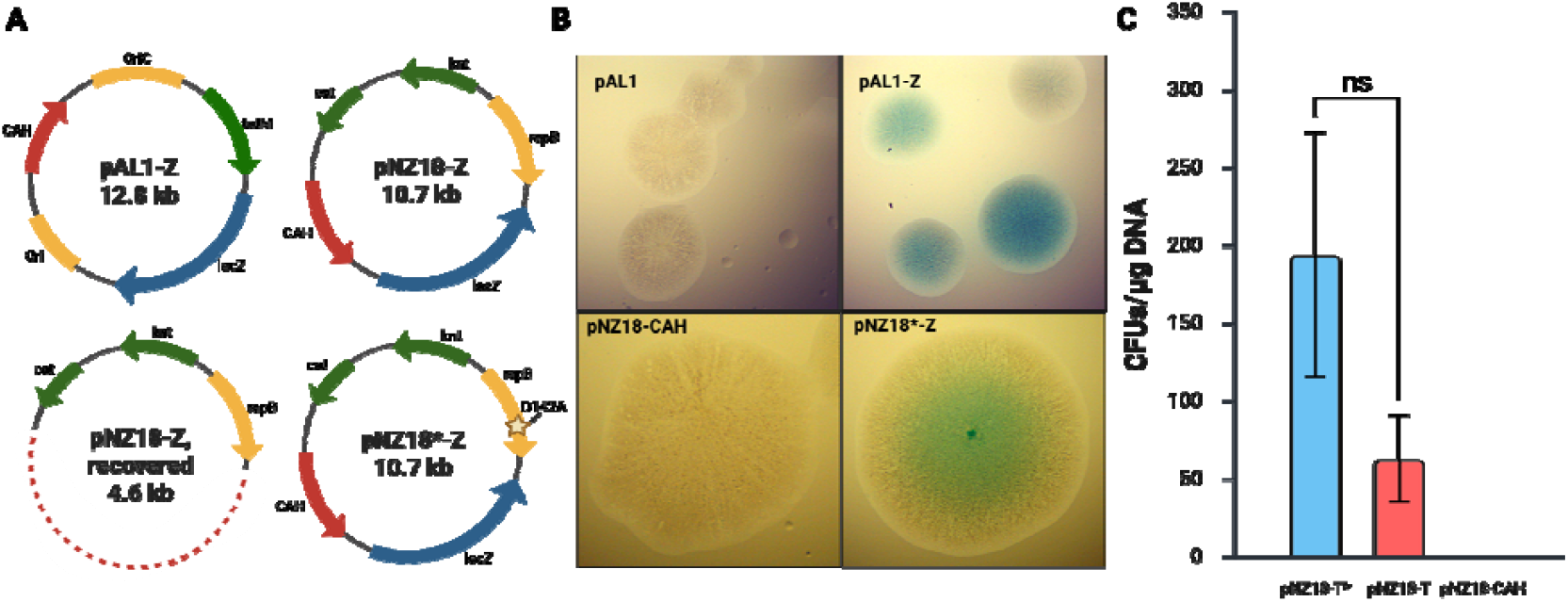
Expresssion of lacZ in pAL1 and pNZ18-CAH. (A) Plasmid maps of pAL1-Z, pNZ18-Z before and after *A. laidlawii* transformation, and the mutant plasmid pNZ18*-Z. OriC = *A. laidlawii* origin of replication, Ori = *E. coli* origin of replication, tetM = tetracycline-resistance gene. Cat = chloramphenicol resistance gene, knt = kanamycin/neomycin resistance gene, CAH = yeast elements (CEN6, ARSH4, HIS3). The dash line in recovered pNZ18-Z shows the portion of the plasmid that was deleted. (B) Colonies of *A. laidlawii* strains carrying plasmids with and without lacZ on selective SP-4 media supplemented with 50 μg/mL X-gal. (C) Transformation efficiency of pNZ18-T and pNZ18*-T to *A. laidlawii*. Transformants are plated on SP-4 media supplemented with both neomycin and tetracycline. Bars show an average from three independent experiments; error bars show standard error of the mean. As a control, transformation efficiency of pNZ18-CAH, which does not ha e a tetracycline-resistance gene, is included. Differences were considered not statistically significant (n.s.) for p > 0.05.

We submitted one recovered pNZ18-Z clone for whole-plasmid sequencing and discovered a large deletion spanning from the yeast elements to the lacZ cassette, resulting in a size reduction from 10.7 kb to 4.6 kb. The replicative elements of pNZ18 are from the rolling-circle replication (RCR) *Lactococcus lactis* plasmid pSH71^28^; RCR plasmids are known to sometimes be segregationally and structurally unstable^29–31^. Indeed, similar large-scale deletions have been reported when transforming strain 8195 with two streptococcal plasmids that are also thought to replicate via RCR^32^. Increasing sizes of heterologous inserts in RCR plasmids is known to promote the formation of high-molecular-weight (HMW) plasmid multimers, which may impact plasmid stability^29,31,33^. In our previous work, and in the work of others using pNZ18-related derivatives, this plasmid was reported to be structurally stable^19,26^. Thus, stability issues may only arise when cloning larger sequences, such as the 3-kb lacZ gene, especially considering that pNZ18-CAH already contains additional sequences for maintenance and selection in yeast (∼ 1.5 kb).

We retested transformation of *A. laidlawii* with pNZ18-Z, this time using DNA from an initial pool of approximately 40 lacZ-positive *E. coli* clones. Like before, we observed white transformants on X-gal plates, but we also identified a single blue *A. laidlawii* colony. This transformant was passaged several times on solid media and always produced blue colonies. We sequenced this plasmid and found it to be completely intact with no structural changes. Instead, we identified only one novel mutation, a single-nucleotide variant in the repB gene causing a codon change from aspartic acid to alanine at amino acid position 142. The repB protein is involved in both replication initiation and termination in RCR plasmids^31^. Derivates of pNZ18 containing the repB D142A mutation are denoted as pNZ18*.

### Investigation of pNZ18*

To investigate whether this repB mutation results in increased plasmid stability, we transformed the original plasmid pNZ18-Z and the mutant plasmid pNZ18*-Z to *A. laidlawii*. Surprisingly, transformation of both plasmids yielded only white colonies on X-gal plates. We considered the possibility that the lacZ-positive *A. laidlawii* transformant may originate from a mutation in the strain rather than the plasmid. To test this, we first passaged this strain several times without antibiotic selection to promote curing of pNZ18*-Z. A single cured strain was then transformed with both versions of the plasmid, but we still saw only white colonies on X-gal plates. Transformants were scraped from plates and pooled in liquid cultures; adding X-gal to these cultures still resulted in no visible blue color.

We next transformed each version of the plasmid to *E. coli*, where we saw a substantial difference in the proportion of blue colonies between pNZ18-Z and pNZ18*-Z colonies (Supplementary Figure 1), suggesting that the repB mutation may indeed play some role in increasing the stability of this plasmid.

The stability of pNZ18*-Z and the potential effects of the repB mutation are not well understood, and it is currently unknown why our attempts to replicate lacZ expression in *A. laidlawii* were unsuccessful. The sequence surrounding D142 in repB from pNZ18 has a high similarity to a flexible hinge region identified in the repB protein from the streptococcal plasmid pMV158^34^ (Supplementary Figure 2), which is in the same plasmid family as pSH71^35,36^. This hinge region separates two main domains of the protein and contributes to flexibility that may be important for its function^34^. It could be possible that this mutation lowers the activity of repB, resulting in less frequent replication initiation and leading to a lower plasmid copy number in the cell. In the RCR plasmid pUB110, a mutation that lowered plasmid copy number was shown to improve stability when cloning heterologous genes in *Bacillus subtills*^*37*^.

To investigate whether plasmid instability only occurs when cloning lacZ, we cloned the ∼ 2-kb tetM gene from pAL1 in pNZ18-CAH at the same location where lacZ was cloned, creating pNZ18-T. We then created a second version of the plasmid, pNZ18*-T, that contained the repB D142A mutation. Wild type *A. laidlawii* DN-E was transformed with both plasmids in parallel, and transformants were plated on neomycin-tetracycline double antibiotic selection. Transformants should only survive if the plasmid does not undergo large deletions that would disrupt function of the tetM gene. Colonies appeared for both transformations, and there was no statistically significant difference in transformation efficiency between the two plasmids (Figure 2C). Yet, colony counts were much lower than what we would expect for pNZ18-CAH. Recovery and sequencing of plasmids from 3 pNZ18-T and pNZ18*T transformants revealed no large-scale deletions in any of them. Thus, pNZ18 derivatives carrying tetM may be more stable than those carrying lacZ. The streptococcal tetM gene^38^ is comparable in size to the *E. coli*-derived lacZ gene, but tetM is lower in G/C content (32% vs 56%). If we assume that higher stability correlates to lower HMW multimer formation in our plasmids, then our results would agree with previous observations that such multimer formation in RCR plasmids is more prevalent when cloning sequences derived from *E. coli*^*33*^.

### Complementation of recA in strain DN-E

Now that we have established exogenous-gene expression in *A. laidlawii* using our shuttle vectors, we attempted to complement recA in strain DN-E. Given that the *A. laidlawii* recA gene is small and low in G/C content, we did not expect stability issues when cloning this gene in pNZ18. We PCR-amplified the recA CDS from the PG-8A genome and cloned it in place of the lacZ gene in pNZ18*-Z, where it would be under control of the strong rpsJ promoter. The complementation plasmid was named pNZ18*-C (Figure 3A). We could not assemble or clone this plasmid in *E. coli*. Issues with cloning *A. laidlawii* recA in *E. coli* have been noted previously, and it is possibly due to toxicity from heterologous recA expression^22^. Instead, pNZ18*-C was assembled in *S. cerevisiae*, and the plasmid was directly transferred from yeast to *A. laidlawii*.

**Figure 3.**
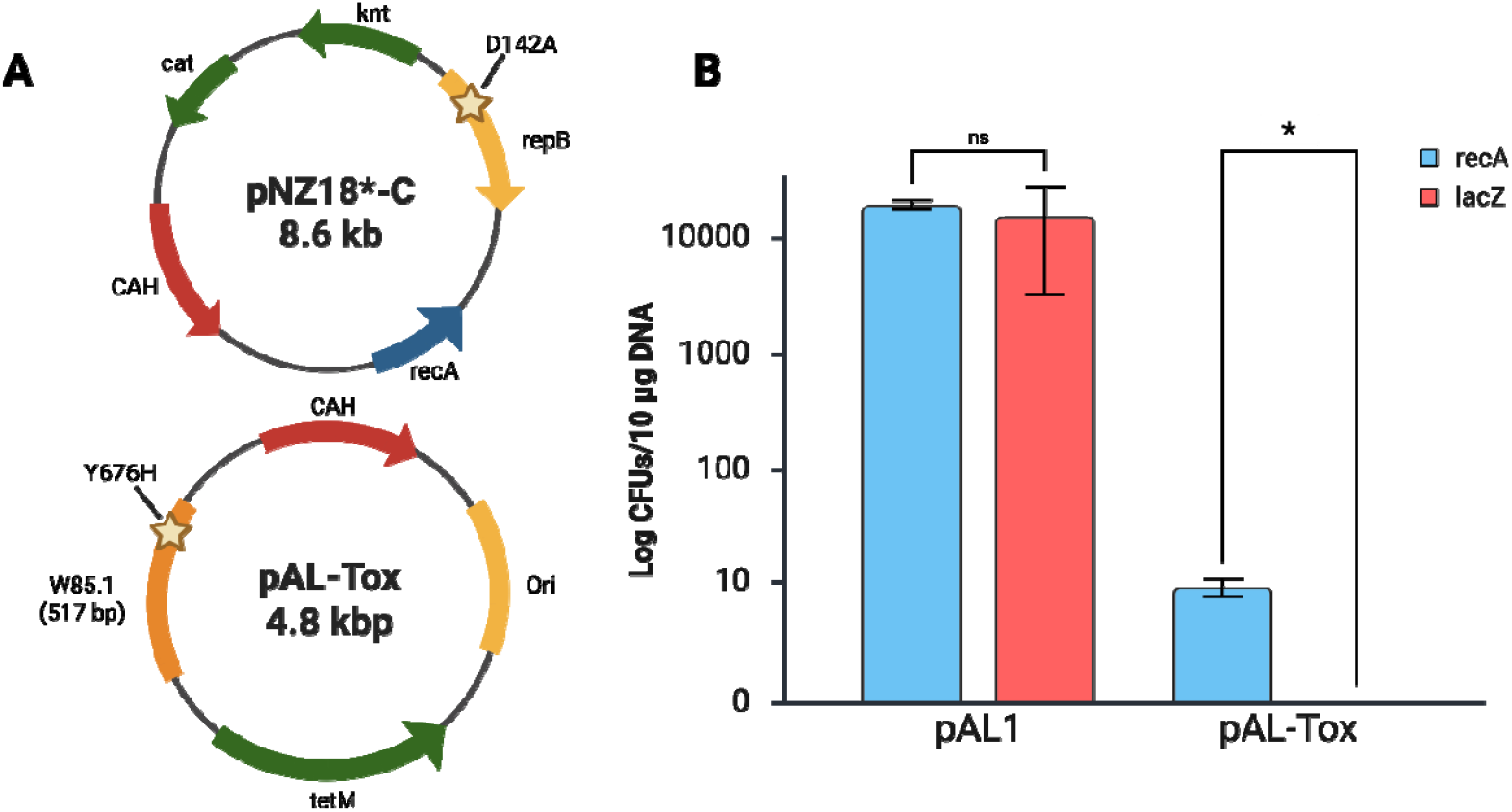
recA complementation of *A. laidlawii* DN-E. (A) Plasmid maps for pNZ18*-C and pAL-Tox. Cat = chloramphenicol resistance gene, knt = kanamycin/neomycin resistance gene, CAH = yeast elements (CEN6, ARSH4, HIS3), tetM = tetracycline resistance gene, Ori = *E. coli* origin of replication. W85.1 = a 517-bp region of homology for the putative toxic gene in the *A. laidlawii* DN-E genome. (B) Comparison of transformation efficiency for pAL1 and pAL-Tox to *A. laidlawii* carrying pNZ18*-C (expressing recA) and pNZ18*-Z (expressing lacZ), standardized to 10 micrograms of plasmid DNA. Bars represent the mean from 3 independent biological replicates, with pAL1 transformations having two technical replicates and pAL-Tox transformations having one technical replicate. Error bars represent standard error of the mean. Preparation of cultures varied slightly between experiments, as described in the Methods section. Statistical significance (p < 0.05) is indicated by an asterisk; n.s. = not significant.

We next constructed an integrative plasmid called pAL-Tox, which contains a region of homology for the last 517 base pairs of the W85.1 CDS. In ACL_0117, this region encodes a conserved endonuclease domain that is the main cause of toxicity when the gene is cloned in yeast and bacteria^18^. Initial attempts to clone pAL-Tox in *E. coli* were unsuccessful, and we suspected that there was some level of expression of the endonuclease domain that was causing toxicity. We were able to mitigate this toxicity by introducing a mutation to a conserved residue, Y676H, which was identified previously^18^. pAL-Tox contains elements for selection and replication in both yeast (CEN6, ARSH4, HIS3) and *E. coli*; the plasmid contains tetM for selection in *A. laidlawii*, but it does not contain a replication origin for this species.

To confirm that our complementation approach was both working and necessary for integration, we transformed pAL-Tox into two separate *A. laidlawii* strains, carrying either pNZ18*-Z or pNZ18*-C. We also transformed each strain with pAL1 as a positive control. As expected, we observed comparable transformation efficiencies of pAL1 between both strains, but we only observed pAL-Tox transformants for the recA-complemented strain (Figure 3B). Transformation to pAL-Tox was notably inefficient; 10 micrograms of plasmid DNA yielded an average of 9 colonies across 3 independent experiments. This reflects an approximate 1000-fold decrease in transformation efficiency compared to transformation with the same mass of pAL1 DNA.

The transformation frequency of a non-replicative plasmid to *A. laidlawii* JA1 was noted to be 10-fold lower compared to previously reported frequencies for replicative and transposon-bearing plasmids in strain 8195^20,39^. This is a much lower difference compared to what we observed. We have noted that strain DN-E shows around 100 times improvement in transformation efficiency of pAL1 compared to strain 8195^19^. It may be possible that the improved-transformation phenotype seen in strain DN-E is specific to replicative plasmids. Alternatively, we have noticed a growth deficit in strains carrying pNZ18*-C, even compared to those carrying pNZ18*-Z. It is possible that expression of recA with a relatively strong promoter is detrimental to cell function, and we may see improved growth and integration rates if we replaced the rpsJ promoter with a weaker one. Including a larger region of homology (>= 1 kb) in future integrative constructs may also improve transformation rates.

### Testing Genomic Integration of pAL-Tox

We performed a series of experiments to confirm that pAL-Tox did indeed integrate into the *A. laidlawii* genome at W85.1. We first isolated DNA from three strains of recA-complemented *A. laidlawii* transformed with pAL1 and three strains transformed with pAL-Tox. Isolated DNA was transformed to *E. coli* to recover plasmids. If pAL-Tox is maintained as an extrachromosomal element, we would expect similar rates of plasmid recovery to pAL1. Across three biological replicates for all 3 strains, recovery of pAL1 yielded an average of 88 *E. coli* colonies per transformation, while recovery of pAL-Tox yielded an average of 0.9 colonies per transformation (Figure 4A).

**Figure 4.**
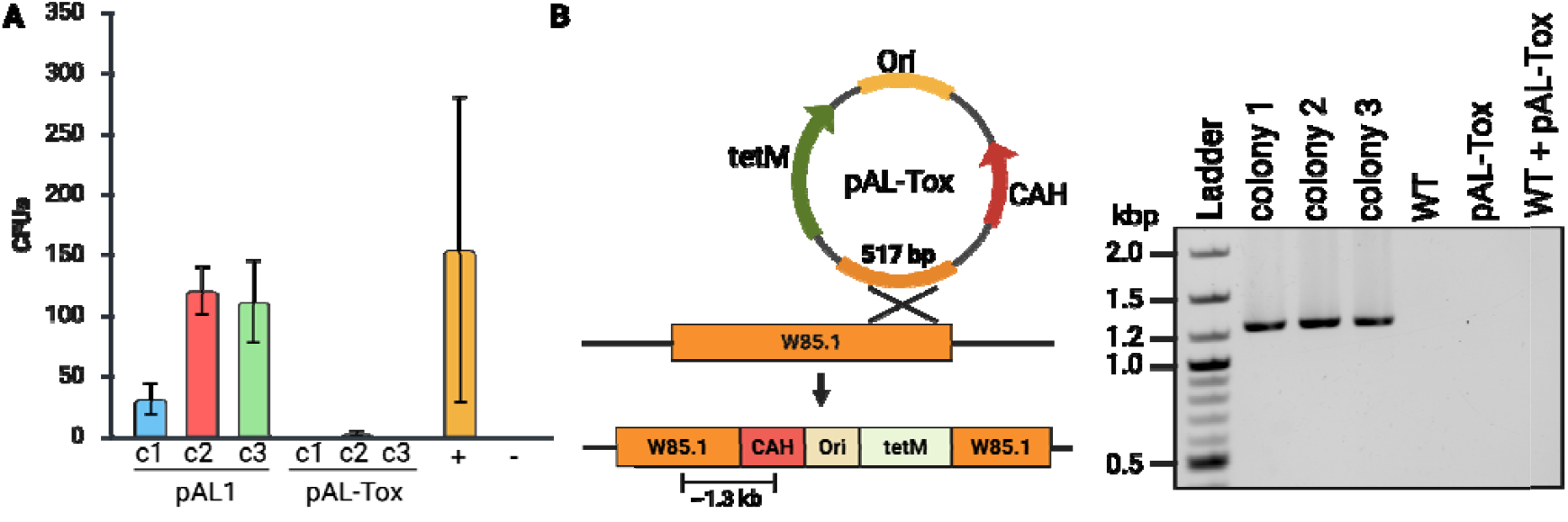
Confirming integration of pAL-Tox in recA-complemented *A. laidlawii*. (A) Recovery of plasmid DNA in *E. coli* from 3 *A. laidlawii* strains transformed with pAL1 or pAL-Tox. Bars represent the mean from 3 independent transformations, with error bars showing the standard error of the mean. The positive control (+) in each experiment was pAL1 isolated from *E. coli*, with colony counts adjusted to 30 nanograms of DNA. The negative control (-) for each experiment was a transformation with no DNA added. (B) Diagram showing expected recombination of pAL-Tox into W85.1. PCR amplifying a junction between the *A. laidlawii* genome and pAL-Tox produces an amplicon of expected size in pAL-Tox transformants, but not for wild type *A. laidlawii* (WT), pAL-Tox plasmid DNA, or a mix of wild type genomic DNA and plasmid DNA (WT + pAL-Tox).

Next, we tested for presence of pAL-Tox in the genome using PCR. We designed primers to amplify a junction between the *A. laidlawii* genome and pAL-Tox based on how this plasmid would recombine into W85.1 (Figure 4B). We observed an expected PCR product of approximately 1.3 kb when we amplified genomic DNA from pAL-Tox transformants. In comparison, we did not see any amplification when we used wild type genomic DNA, purified pAL-Tox DNA, or a mixture of both as PCR templates. These results strongly support that we were successful in integrating pAL-Tox directly into W85.1.

### Integration of pAL1

We were curious if we could also integrate pAL1 into the genome of our recA-complemented strain. OriC-based plasmids developed for other *Mollicutes* are known to recombine into the host origin of replication^40–43^, bu we had not observed this previously in wild type *A. laidlawii* 8195^19^. To this end, we pooled several pNZ18*-C strains transformed with pAL1 in a liquid culture, and this pool was passaged three times with tetracycline and neomycin selection to maintain both plasmids and provide opportunity for pAL1 integration (Supplementary Figure 3A). We passaged the pool another six times without any selection to cure the strains of non-integr ted plasmids. We then did a final passage with tetracycline to select for strains that had retained pAL1 in the genome. We isolated 3 individual strains from this final pool, and we used another PCR screen to test for integration of pAL1 into the OriC. We saw an expected amplicon of approximately 3 kb when using DNA from these strains as template, but no amplification was observed for wild type genomic DNA, isolated pAL1 DNA, or a mixture of both templates (Supplementary Figure 3B).

Alongside the pNZ18*-C pool, we also tested pAL1 integration in two other transformation pools, one with strains carrying a second recA-complementation construct that included the whole recA operon from strain PG-8A (pNZ18*-C2) and one with strains carrying pNZ18*-Z, containing no recA complement. Throughout passaging, aliquots of each culture were taken for DNA isolation and PCR screening. Shown in Supplementary Figure 3C are the results of a PCR screen for all three cultures at passages 1, 3, and 10. The expected amplicon is only seen in recA-complemented pools at the first passage. By the third passage, the amplicon is also seen in the non-complemented pool, but unlike the complemented pools, it is faint and does not increase in intensity by the final passage. We spotted serial dilutions of passage 10 cultures on non-selective and tetracycline-supplemented SP-4 agar plates (Supplementary Figure 3D). While growth across all three pools was comparable on non-selective media, the recA-complemented pools grew to approximately 3 more dilutions than the non-complemented pool on selective media, showing that the non-complemented population had a substantially lower proportion of pAL1-integrated cells.

Although we had previously reported no evidence of pAL1 integration for wild type *A. laidlawii*, we have seen here that some proportion of cells lacking recA complementation still show integration into the OriC after several generations. It is possible that the truncated recA protein in wild type cells supports low levels of recombination, or that integration of pAL1 is occurring through a recA-independent mechanism. As assessed through our PCR screen, there was very little or no integration in non-complemented cells before the first passage. Thus, recA complementation appears necessary for integration of non-replicative or otherwise transiently maintained plasmids, which is consistent with what we observed for transformation of pAL-Tox.

Despite the PG-8A recA gene in pNZ18*-C having a high sequence homology to the DN-E recA gene, all the final strains we generated were sensitive to neomycin, suggesting that the plasmid had been lost and not integrated into the genome. pNZ18 is derived from the high-copy-number plasmid pNZ12^28^. It is possible that the level of neomycin resistance conferred by the knt marker is not sufficient for antibiotic selection when the plasmid is present in the genome as a single copy. Our complementation plasmid thus presents a useful way to “toggle” homologous recombination in this strain, where it can be transformed into cells to enable integration and then lost through plasmid curing.

### Genome Transfer to S. cerevisiae

Finally, we planned to transfer the genome of one pAL-Tox integration strain to *S. cerevisiae* using cell fusion. We expected that cloning of the whole genome would only be possible when W85.1 was disrupted. To test this, we also performed cell fusion with one pAL1 integration strain; pAL1 contains the same YCp elements as pAL-Tox, but because W85.1 is uninterrupted in this strain, we only expected partial genome cloning to occur (Figure 5A). Saturated cultures of both strains were concentrated and transformed into *S. cerevisiae* spheroplasts, and the yeast were recovered and plated on synthetic media lacking histidine. We isolated DNA from 20 yeast transformants from each cell fusion experiment and performed multiplex PCR to screen for the presence of 3 loci across the DN-E genome. We found that 13/20 yeast colonies screened positive for all 3 loci when cell fusion was performed with the pAL-Tox integration strain, while 0/20 colonies screened positive for all 3 loci when cell fusion was performed with the pAL1 integration strain (Supplementary Figure 4).

**Figure 5.**
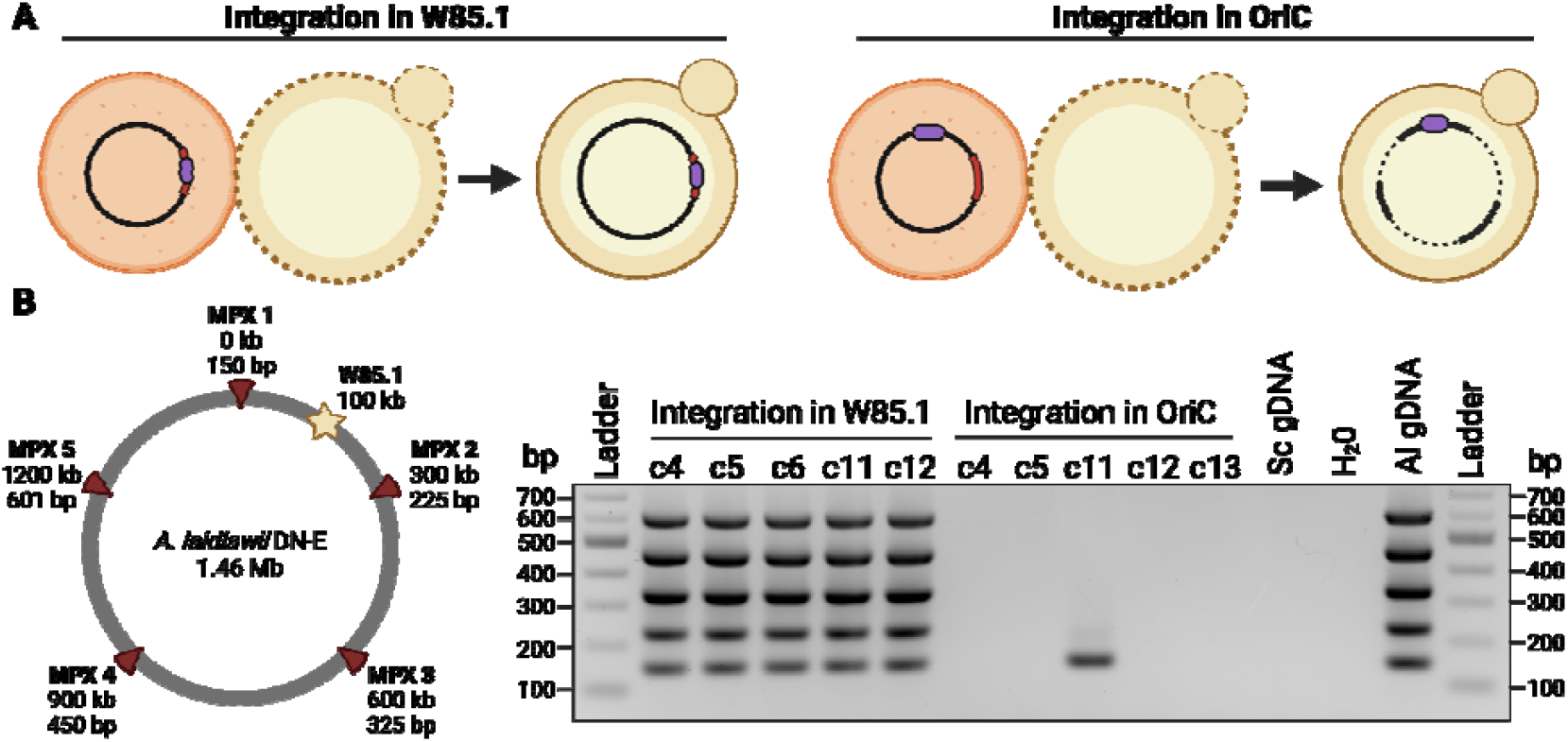
Cloning the *A. laidlawii* DN-E genome in yeast using cell fusion. (A) Expected outcome for cell fusion to yeast using *A. laidlawii* strains with yeast elements (represented as a purple oval) integrated into the putative toxic gene W85.1 (represented as a red region in the genome) or into the origin of replication (OriC). (B) Screening yeast colonies for presence of the entire *A. laidlawii* genome with multiplex PCR. A legend shows the approximate location and amplicon size of each primer pair; the approximate location of the toxic gene W85.1 is also shown. Controls included using yeast (Sc) genomic DNA, sterile water, and *A. laidlawii* (Al) genomic DNA.

We then selected 5 of the initially screened 20 yeast colonies from each cell fusion experiment for a higher-resolution multiplex screen, amplifying every 300 kb in the 1.46 Mb genome for a total of 5 loci. 5/5 strains that were fused with pAL-Tox-integrated *A. laidlawii* tested positive for the 5 loci, whereas 0/5 strains fused with pAL1-integrated cells tested positive for all 5 loci (Figure 5B). One of the clones in the latter group did test positive for a single locus, suggesting that each of these yeast clones may carry some fragmented or incomplete portion of the genome. This seems likely, considering that these strains must retain the HIS3 gene to grow on selective media.

Given these results, and the nature of the pAL-Tox integration, we expect that the function of W85.1 in transformed *A. laidlawii* is either partially or completely disrupted, which would suggest that this gene is not essential. ACL_0117 is predicted to be an extracellular nuclease in strain PG-8A^18^. We have observed rapid digestion of extracellular plasmid DNA for wild type *A. laidlawii* PG-8A and 8195^19^; it is possible that W85.1 encodes a nuclease that contributes to this digestion. Thus, future experiments should be conducted to test whether pAL-Tox strains show improved transformation rates. This may be an important consideration for PEG-mediated transformation, where cells and DNA are incubated together for longer than in electroporation.

In conclusion, we have shown that complementation of *A. laidlawii* strain DN-E with recA enables targeted genomic integration using a non-replicative plasmid. Inactivation of W85.1 in strain DN-E was necessary for cloning the genome in yeast, which agrees with past work done in strain PG-8A. Delivery of *Acholeplasma* genomes to yeast using cell fusion is a significant improvement to the TAR cloning approach that was used previously, as it is more efficient and does not require the isolation of large, delicate genomes. Future work will focus on developing genome transplantation to *A. laidlawii* to deliver such yeast-cloned genomes back to recipient bacteria, which is the final step in our goal to create synthetic *Acholeplasma* strains.

## MATERIALS AND METHODS

### Strains and Cultures

*A. laidlawii* strain DN-E (ATCC: BAA-3342) was grown at 32°C in SP-4 media lacking serum^19^ without shaking. For antibiotic selection, SP-4 was supplemented with 100 μg/mL neomycin and/or 1 μg/mL tetracycline. Solid media was made with 1% agar.

*E. coli* strains MC1061 and Epi300 were grown in LB media (for 1 liter: 10 g tryptone, 10 g sodium chloride, 5 g yeast extract) at 37°C with shaking at 225 rpm. Solid media was made with 1.5% agar. For antibiotic selection, media was supplemented with 10 μg/mL chloramphenicol and/or 10 μg/mL tetracycline. For both *A. laidlawii* and *E. coli*, plates were supplemented with 50 μg/mL X-gal for blue-white screening.

*S. cerevisiae* strain VL6-48 was grown in 2x YPAD broth (for 1 liter: 20 g yeast extract, 40 g peptone, 20 g D-glucose, 160 mg adenine hemisulfate) at 30°C with shaking at 225 rpm. For transformation, cells were instead grown in minimal synthetic defined media with histidine dropout supplement (Takara Bio) and 1 M D-sorbitol. Solid media was made with 2% agar.

### Transformation to *A. laidlawii*

*A. laidlawii* was transformed as described previously^19^. Cultures were started by inoculating 5 mL of SP-4 (supplemented with appropriate antibiotics) with an aliquot of cells from a -80°C stock. After 3 days of growth at 32°C, cultures were passaged 1/5 in 10 mL of fresh SP-4 and grown overnight. The next day, 1-mL aliquots were centrifuged at 15,000 rpm for 10 minutes at room temperature. Cells were washed with an equivalent volume of sucrose-HEPES buffer (272 mM Sucrose, 8 mM HEPES, pH = 7.4) and centrifuged as before. Cells were then resuspended in 100 μL of the same buffer and kept on ice for 5 minutes. Plasmid DNA dissolved in ddH^2^O was added to cells, and the mixture was kept on ice for 2–3 minutes. Next, the mixture was transferred to a pre-chilled 2 mm cuvette (VWR) and pulsed at 2.5 kV for 5 ms (200 Ω, 25 μF) in a Bio-Rad GenePulser Xcell. One milliliter of cold SP-4 was added, and the cuvettes were kept on ice for 10 minutes. Cells were then transferred to 1.5 mL tubes and recovered at 32°C for 2 hours. Between 100 and 250 μL of recovered cells were plated on one SP-4 plate, supplemented with appropriate antibiotics.

Three independent experiments were performed to test transformation of pAL1 and pAL-Tox to *A. laidlawii* strains carrying pNZ18*-C or pNZ18*-Z. In cases where the growth of pNZ18*-C cultures was visibly slower than pNZ18*-Z cultures, dilutions were made to account for this difference in growth rate. For all experiments, pNZ18*-C and pNZ18*-Z cultures were started in 5 mL of SP-4 with neomycin and grown for three days at 32°C. On the third day, pNZ18*-Z cultures for Experiments 1 and 3 were passaged 1/2 in 6 mL SP-4 and grown overnight alongside undiluted pNZ18*-C cultures. The next day, both cultures were passaged 1/5 in fresh media and incubated overnight prior to transformation the next day. In Experiment 2, the intermediate dilution of pNZ18*-Z was omitted, and both cultures were passaged 1/5 in fresh media after the third day of growth prior to transformation the next day. For each experiment, culture growth was found to be similar between both strains as visually assessed by bacterial pellet size and phenol red color change.

### Transformation to *E. coli*

*E. coli* was grown in 5 mL of LB overnight from a single colony. The next day, the culture was diluted 1/100 in 50–100 mL LB and grown to OD_600_ = 0.5–0.7. Cultures were transferred to a 50 mL conical tube and centrifuged at 3,000 x g for 15 minutes at 4°C. For preparation of electrocompetent *E. coli*, cells were washed three times in an equivalent volume of ice-cold 10% glycerol and resuspended in 1/100^th^ the initial volume with 10% glycerol. For preparation of chemically competent, we followed a protocol from Alice Pawlowski^44^ with slight modifications. *E. coli*, cells were resuspended in an equivalent volume of ice-cold 0.1 M CaCl_2_ and incubated on ice for 1 hour. The cells were centrifuged as before and resuspended in 1/100^th^ the initial volume with 0.1 M CaCl_2_, 15% glycerol. Both preparations of cells were stored at -80°C until use.

Each *E. coli* transformation used 25–30 μL of thawed competent cells. For electroporation, cells were pulsed using the same cuvettes and conditions as *A. laidlawii*. For chemical transformation, competent cells were mixed with plasmid DNA and incubated on ice for 30 minutes. Cells were transferred to a 42°C water bath for 45 seconds and put back to ice for 5 minutes. Following each transformation, cells were recovered with 450 μL SOC media (For 1L: 20 g tryptone, 5 g yeast extract, 0.5 g NaCl, 5 mL 2M MgCl, 10 mL 250 mM KCl, 20 mL 1 M glucose) for 1 hour at 37°C with shaking at 225 rpm. Recovered cells were centrifuged at 10,000 rpm for 5 minutes at room temperature, and the pellet was resuspended in 100 μL LB for plating.

For transformation of *E. coli* with pNZ18-Z and pNZ18*-Z, 100 ng of plasmid DNA was electroporated to competent MC1061 cells, and a negative control was included where no DNA was added. Centrifugation of recovered cells was omitted, and 100 μL of transformed cells were plated on LB agar plates supplemented with chloramphenicol and X-gal.

### Plasmid recovery from A. laidlawii to E. coli

*A. laidlawii* transformants were passaged on selective solid SP-4 media. Streaks were then inoculated to 5 mL of selective liquid culture with a P1000 pipette tip and grown for 4–7 days at 32°C. 1-milliliter aliquots of densely grown culture were used for alkaline lysis using Buffers P1, P2, P3 (Qiagen) with alcohol precipitation, and the resulting DNA was resuspended in 50 μL ddH_2_O.

For comparing plasmid recovery between pAL1 and pAL-Tox *A. laidlawii* transformants, 20 μL of isolated DNA was used for electroporation to *E. coli* strain Epi300 as described above, where 30–100 ng of pAL1 DNA from *E. coli* was used as a positive control, and a transformation with no DNA was used as a negative control. For recovery of pNZ18-T and pNZ18*-T plasmids, 2 μL of isolated *A. laidlawii* DNA was transferred to *E. coli* strain MC1061 using chemical transformation. Transformants were recovered on chloramphenicol-tetracycline LB media, and one colony from each transformation was sent for sequencing.

### Plasmid Sequencing

Plasmids were isolated from *E. coli* using the EZ-10 Spin Column DNA Cleanup Miniprep Kit (BioBasic) or the Monarch Plasmid Miniprep Kit (NEB). Miniprepped samples were submitted to Plasmidsaurus (plasmidsaurus.com) for whole-plasmid sequencing.

### Cell Fusion

Yeast cells were grown, spheroplasted, and prepared for cell fusion as described previously^3^. 50 mL of densely grown *A. laidlawii* cultures were treated with 100 μg/mL chloramphenicol and incubated at 32°C for 1 hour. Cultures were centrifuged at 3,000 x g at 4°C for 20 minutes, and the resulting cell pellets were resuspended in 100–500 μL of resuspension buffer (0.5 M sucrose, 10 mM Tris-HCl [pH = 8], 10 mM CaCl_2_, 2.25 mM MgCl_2_, pH adjusted to 8) followed by heat inactivation at 49°C for 10 minutes. 50 microliters of resuspended *A. laidlawii* cells were mixed with 200 μL of yeast spheroplasts. Following recovery, yeast spheroplasts were mixed with 5– 8 mL of molten -HIS agar and plated on 30 mL of previously hardened -HIS agar. Plates were incubated at 30°C for 5–6 days. The resulting yeast transformants were passaged twice on -HIS solid media prior to genotyping with an initial multiplex PCR screen. A subset of colonies was passaged a third time on solid media and then genotyped using a higher-resolution multiplex screen.

## ACKNOWLEDGEMENTS

This work was funded by the Natural Sciences and Engineering Research Council of Canada (NSERC) through project code RGPIN-2025-05428 awarded to B.J.K.

We thank Emma J. Walker for creating the *A. laidlawii* cell illustration used in many of the figures. All figures, including the cover art, were created using Biorender.com.

## AUTHOR CONTRIBUTIONS

Conceptualization: DPN and BJK; Data curation: DPN; Formal analysis: DPN, BJK; Funding acquisition: BJK; Investigation: DPN, TSS; Methodology: DPN, BJK; Project administration: BJK; Resources: BJK, Supervision: DPN, BJK; Validation: DPN, TSS; Visualization: DPN; Writing: DPN; Writing – review & editing: DPN, BJK.

