## Supplementary file for "Direct Genome Transfer from *Acholeplasma laidlawii* to Yeast"

**Supplementary Figures**

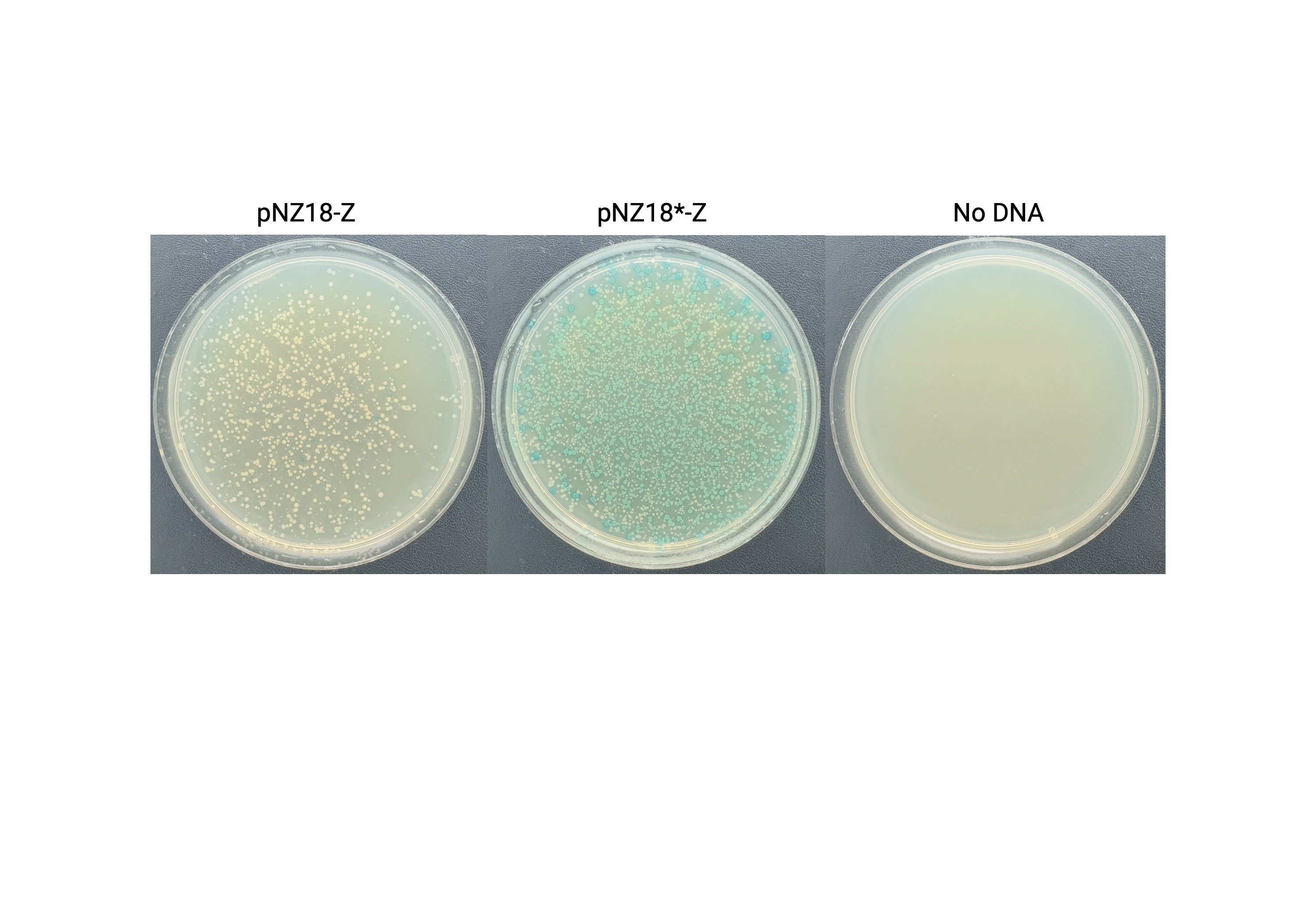
**Supplementary Figure 1. Transformation of wild type and mutant pNZ18-Z to *E. coli*.** Transformation to *E. coli* with pNZ18-Z (wild type), pNZ18*-Z (repB mutation D142A), and no DNA on LB-agar plates supplemented with 10 μg/mL chloramphenicol, 50 μg/mL X-gal.

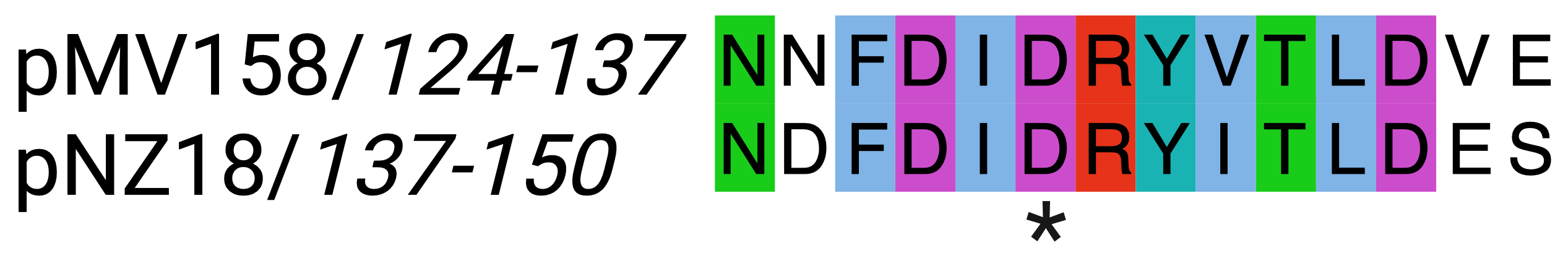

**Supplementary Figure 2. Homology between repB in pMV158 and pNZ18.** A local alignment between the hinge region of pMV158 repB protein (residues 124 – 137; described previously^1^) and a homologous region in pNZ18 (residues 137 – 150), as determined by a global alignment. The aspartic acid residue that is mutated to alanine in pNZ18* is indicated by an asterisk.

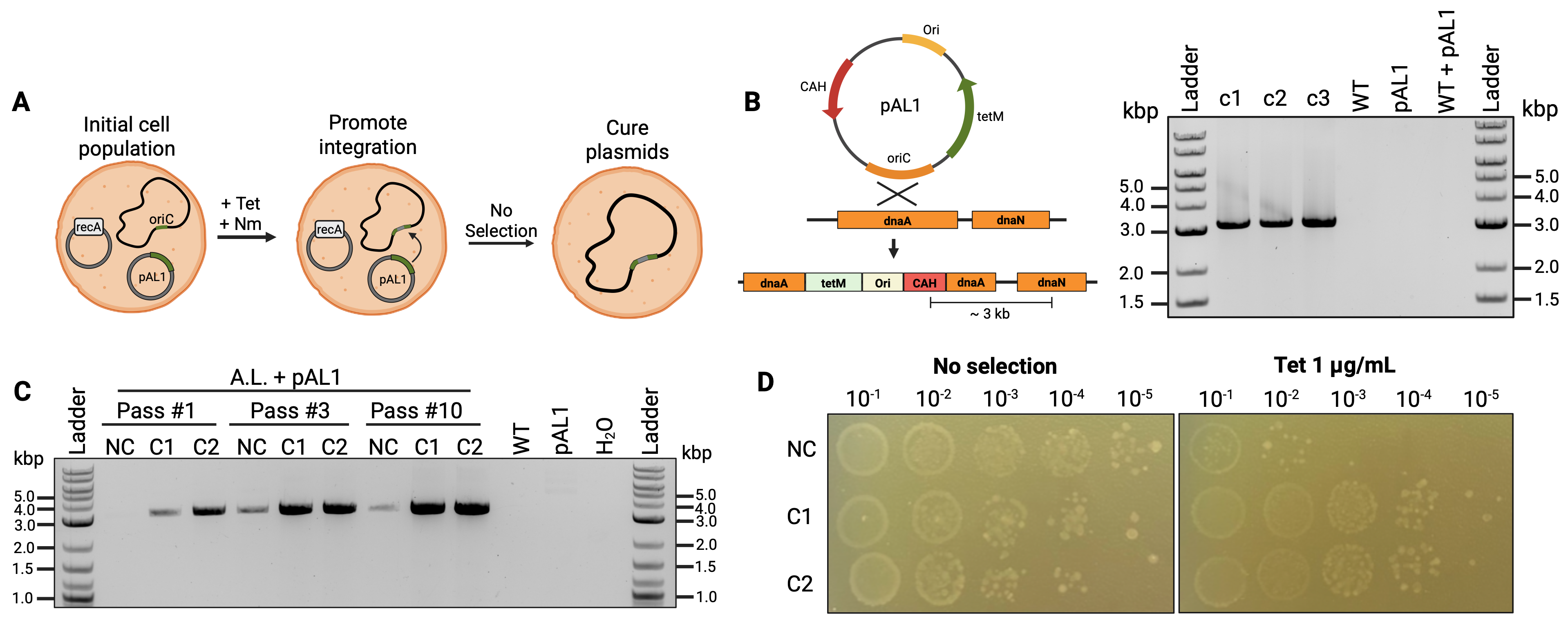

**Supplementary Figure 3. Integration of pAL1 in *A. laidlawii*.** (A) Diagram showing the strategy used to select for pAL1 integration strains following transformation of the plasmid. (B) PCR screening a junction between the *A. laidlawii* genome and pAL1. The expected amplicon is seen for 3 pAL1 integration strains, but not for wild type *A. laidlawii* (WT), pAL1 DNA, or a mix of these templates (WT + pAL1). (C) PCR screening for integration of pAL1 in pools of pAL1-transformed *A. laidlawii* carrying pNZ18*-Z (NC; not complemented), pNZ18*-C (C1), or a second recA-complementation construct pNZ18*-C2 (C2). DNA from passage 1, 3, and 10 cultures was genotyped. (D) Serial dilutions of *A. laidlawii* pools from passage 10, spotted on non-selective SP-4 agar or SP-4 agar supplemented with 1 μg/mL tetracycline. Created using Biorender.com.

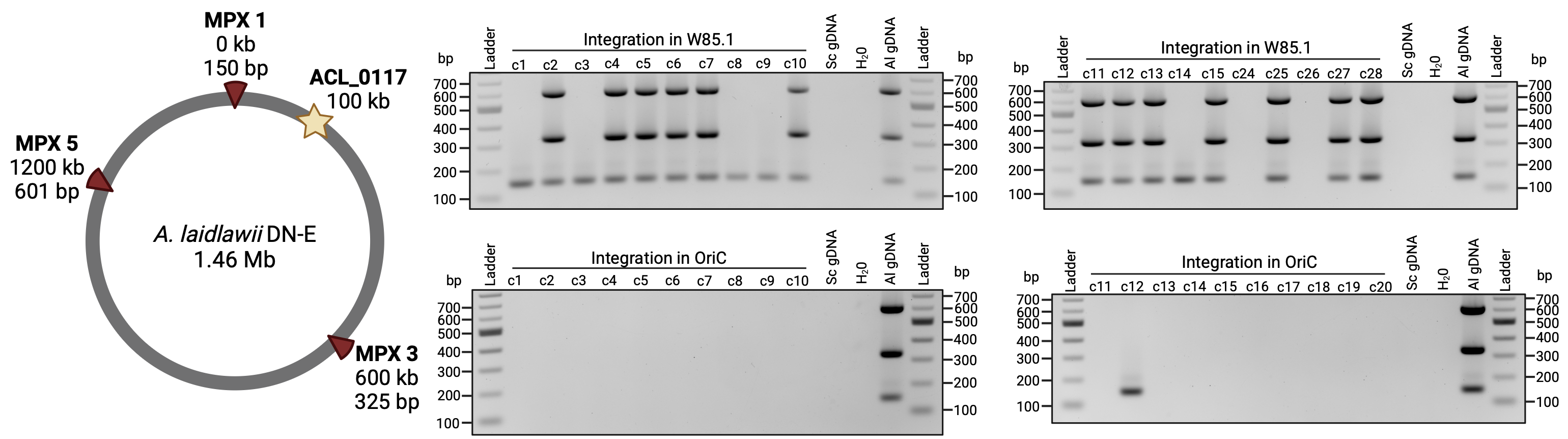

**Supplementary Figure 4. Primary screen of yeast colonies following cell fusion**. Results of a multiplex PCR screen for 20 yeast transformants following cell fusion with pAL-Tox (YCp integration in W85.1) and pAL1 (YCp integration in oriC) *A. laidlawii* strains. Locations of the 3 loci tested are shown on a genome map of strain DN-E. Created using Biorender.com.

**Supplementary Tables**

**Supplementary Table 1.** List of primers used for plasmid assembly. Each primer has a 20-base-pair portion that binds to the template and a 40-base-pair portion that adds homology for an adjacent fragment, the latter of which is indicated by an underline.

| Name | Sequence (5’ – 3’) | Description |
| --- | --- | --- |
| BK3402_F | cacactggctcaccttcgggtgggcctttctgcgtttatagggcccgatcgccaacaaat | Linearizes pAL1 for pAL1-Z |
| BK3402_R | gtacaaagtctggattatatagatcttttcccatcgcatagctactagtattattatgtg | Linearizes pAL1 for pAL1-Z |
| BK2559_F | cgttatatgttcaacaaaatcacataataatactagtagctatgcgatgggaaaagatct | Amplifies PG-8A rpsJ promoter for pAL1-Z |
| BK3041_R | tttgcgtattgggcgccagggtggtttttcttttcaccatttttattctcctttcgccta | Amplifies PG-8A rpsJ promoter for pAL1-Z |
| BK3041_F | taaagtctatgcaaggattataggcgaaaggagaataaaaatggtgaaaagaaaaaccac | Amplifies promoter-less lacZ with *E. coli* terminator for pAL1-Z |
| BK3042_F | agagcaaggtaaaaggtagtatttgttggcgatcgggccctataaacgcagaaaggccca | Amplifies promoter-less lacZ with *E. coli* terminator for pAL1-Z |
| BK2705_F | cacactggctcaccttcgggtgggcctttctgcgtttatagttcattcagggcaccggag | Linearizes pNZ18-CAH for pNZ18-Z |
| BK3158_R | tcaagtccagactcctgtgtaaaactacataagaacacct | Linearizes pNZ18-CAH for pNZ18-Z |
| BK3157_F | aggtgttcttatgtagttttacacaggagtctggacttgatatgcgatgggaaaagatct | Amplifies lacZ construct from pAL-Z for pNZ18-Z |
| BK2706_R | agatggattgcacgcaggttctccggtgccctgaatgaactataaacgcagaaaggccca | Amplifies lacZ construct from pAL-Z for pNZ18-Z |
| BK3115_F | AGGTGTTCTTATGTAGTTTTACACAGGAGTCTGGACTTGAaaacactataaattcgtaat | Amplifies putative ACL_0241 promoter from PG-8A genome for pNZ18-T |
| BK3115_R | CATCAACGTGAGCTAATACACCGATATTTATGATTTTCATattaaaaacctcctacagtt | Amplifies putative ACL_0241 promoter from PG-8A genome for pNZ18-T |
| BK3116_F | TATTGATAAAGTTCGTTATATGTTCAACAAAATCACATAAtaaaaattatctcctctata | Amplifies putative ACL_0241 terminator from PG-8A genome for pNZ18-T |
| BK3116_R | agatggattgcacgcaggttctccggtgccctgaatgaacttatgcatatttttaacctc | Amplifies putative ACL_0241 terminator from PG-8A genome for pNZ18-T |
| BK3117_F | ttaactgtacaaattaggaaaactgtaggaggtttttaatATGAAAATCATAAATATCGG | Amplifies tetM CDS for pNZ18-T |
| BK3117_R | aatgcaaaaaaaatcccctctatagaggagataatttttaTTATGTGATTTTGTTGAACA | Amplifies tetM CDS for pNZ18-T |
| BK3102_F | gttcattcagggcaccggagaacctgcgtgcaatccatct | Linearizes pNZ18-CAH for pNZ18-T |
| BK3158_R | tcaagtccagactcctgtgtaaaactacataagaacacct | Linearizes pNZ18-CAH for pNZ18-T |
| BK3625_F | tgcaaaggttcttgatgctgaaacgggggaaataaaatgacaaacaaagaaaaagagtta | Linearizes pNZ18-T to replace repB |
| BK3625_R | aaaatccaaaatttctagctttagtatttttaatagccatgatatattaccttatcaaaa | Linearizes pNZ18-T to replace repB |
| BK3626_F | tacgagttttcgctacttgtttttgataaggtaatatatcatggctattaaaaatactaa | Amplifies mutant repB gene from pNZ18*-Z |
| BK3626_R | aattcctcattttcagcaaataactctttttctttgtttgtcattttatttcccccgttt | Amplifies mutant repB gene from pNZ18*-Z |
| BK3177_F | taaagtctatgcaaggattataggcgaaaggagaataaaaatgagcgataataaaaaaca | Amplifies recA gene from strain PG-8A for pNZ18*-C |
| BK3176_R | cgttttatttgatgcctggctctagtagcgatctacactatgatagtttcaccactttgt | Amplifies recA gene from strain PG-8A for pNZ18*-C |
| BK3182_F | tagtgtagatcgctactagagccaggcatcaaataaaacg | Linearizes pNZ18*-Z for pNZ18*-C |
| BK3159_R | ttttattctcctttcgcctataatccttgcatagacttta | Linearizes pNZ18*-Z for pNZ18*-C |
| BK3409_F | ttgcaagcagcagattacgcgcagaaaaaaaggatctcaagctactagtattattatgtg | Amplifies tetM for pAL-Tox |
| BK3408_R | ccaataacgaacaatttcatggccagagtaaataccatcttccctttagtgagggttaat | Amplifies tetM for pAL-Tox |
| BK2511_F | aggtgttcttatgtagttttacacaggagtctggacttgatttccataggctccgccccc | Amplifies *E. coli* ori for pAL-Tox |
| BK3409_R | cgttatatgttcaacaaaatcacataataatactagtagcttgagatcctttttttctgc | Amplifies *E. coli* ori for pAL-Tox |
| BK3373_F | tatagattttccggagtatgtagatgtctactttaactagatcacgtgctataaaaataa | Amplifies yeast elements, CEN/ARS/HIS for pAL-Tox |
| BK2511_R | atttttgtgatgctcgtcaggggggcggagcctatggaaatcaagtccagactcctgtgt | Amplifies yeast elements, CEN/ARS/HIS for pAL-Tox |
| BK3408_F | agggttttcccagtcacgacattaaccctcactaaagggaagatggtatttactctggcc | Amplifies 517 base pairs of W85.1 for pAL-Tox |
| BK3401_R | ttaaaaaatttaaattataattatttttatagcacgtgatctagttaaagtagacatcta | Amplifies 517 base pairs of W85.1 for pAL-Tox |

**Supplementary Table 2.** Primers for testing integration of pAL1 and pAL-Tox in the *A. laidlawii* genome.

| Name | Sequence (5’ – 3’) | Description |
| --- | --- | --- |
| BK3361_F | caccaatacgccatacttgca | Screening integration of pAL-Tox; binds within the 5’ region of ACL_0117 (*A. laidlawii* genome) |
| BK3376_R | gagagcaggaagagcaaggt | Screening integration of pAL-Tox; binds within ARSH4 (plasmid) |
| BK3379_F | cggttgccataagagaagcc | Screening integration of pAL1; binds within HIS3 (plasmid) |
| BK3365_R | acgcccaggtaaagcaattt | Screening integration of pAL1; binds within dnaN (*A. laidlawii* genome) |

**Supplementary Table 3.** Multiplex PCR primers used to amplify loci in the *A. laidlawii* DN-E genome.

| Name | Sequence (5’ – 3’) |
| --- | --- |
| BK2481_F (MPX1_F)  150 bp | atgctgcaggattggtctttg |
| BK2481_R (MPX1_R)  150 bp | acatcatccattcaactgcact |
| BK2482_F (MPX2_F)  225 bp | attgatcatgtaggtattgggct |
| BK2482_R (MPX2_R)  225 bp | ttcctttctttctaatcaggcgt |
| BK2497_F (MPX3_F)  325 bp | ttagagcctacagcgccaat |
| BK2497_R (MPX3_R)  325 bp | agacacccgattcccattta |
| BK2498_F (MPX4_F)  450 bp | aaagcatggatagacaaaatagagc |
| BK2498_R (MPX4_R)  450 bp | cggtagcaaatcaattgaaaca |
| BK2485_F (MPX5_F)  601 bp | aagtcgataaaggcatgatgtgt |
| BK2485_R (MPX5_R)  601 bp | cacttgttggtgcgatggta |

ADDITONAL MATERIALS AND METHODS

**pAL1 Integration**

Three *A. laidlawii* strains were transformed with pAL1, carrying either pNZ18*-Z or one of two recA-complementation plasmids: pNZ18*-C, or a second recA complementation carrying the PG-8A recA operon, pNZ18*-C2. Approximately 100–300 colonies from each transformation were scraped from agar plates and separately inoculated to 10 mL of SP-4 media supplemented with tetracycline and neomycin. The cultures were passaged 3 more times in 10 mL SP-4 media with neomycin and tetracycline; the dilution factor of each passage ranged from 100 to 1,000. The cultures were then passaged 1/1,000 once in 5–10 mL SP-4 with tetracycline.

Next, to cure the population of all plasmids, the cultures were passaged 6 times in 5–10 mL of SP-4 without any antibiotics, with a dilution factor ranging from 1,000 to 10,000. To select for cells that retained a genomic copy of pAL1, the cultures were passaged 1/100 in SP-4 media with tetracycline. Once grown, serial dilutions were plated on solid SP-4 media with tetracycline, and individual colonies were screened for genomic integration of pAL1. One milliliter of culture from passages 1, 3, and 10 was saved for DNA isolation and genotyping. Passage 10 cultures were serially diluted, and five microliters of each dilution was spotted on both non-selective and tetracycline-supplemented SP-4 agar.

**repB Sequence Alignment**

The full-length amino acid sequences of repB from pNZ18 and repB from pMV158 (UnitProt ID: P13921)^2^ were first globally aligned using BLASTp^3^. This alignment confirmed homology between the two proteins, with 50.70% total sequence identity. The previously identified 14-amino-acid sequence comprising the hinge region of repB from pMV158^1^ was then locally aligned to the corresponding region of repB from pNZ18 using the EMBL-EBI Clustal Omega tool^4^. The local alignment data was visualized in Jalview^5^ using the Clustal color scheme.

**Plasmid Construction**

Plasmids were assembled through homologous recombination in yeast. Template DNA was PCR amplified with primers to add overlapping homologous sequences to adjacent fragments. PCR primers used for plasmid assembly are listed in Supplementary Table 2. All amplicons were mixed in roughly equimolar amounts and co-transformed to yeast spheroplasts; preparation of spheroplasts is described previously^6^. The resulting yeast transformants were pooled and treated with zymolyase (BioShop) before DNA isolation with alkaline lysis using buffers P1, P2, and P3 (Qiagen), followed by alcohol precipitation. Isolated DNA was transformed to *E. coli*, and individual clones were screened and sequenced for correct plasmid assembly.

The plasmid pAL1-Z is derived from a reduced version of pAL1, where approximately 1.6 kb–including a truncated lacZ gene, a puromycin resistance gene, and a homologous sequence for the gene downstream of ACL_0117–was excluded. A 368-base-pair fragment containing the putative promoter sequence for the rpsJ (ACL_0086) operon was amplified from *A. laidlawii* PG-8A genomic DNA. The full-length lacZ CDS and double *E. coli* terminator (iGEM Part BBa_B0015) were amplified from an unpublished plasmid generously provided by Dr. Gregory Pellegrino (The University of Western Ontario, Department of Biochemistry). pNZ18-Z was constructed by amplifying the new lacZ cassette from pAL1-Z and cloning it into pNZ18-CAH, which was first linearized by PCR.

For pAL-Tox, YCp elements (CEN6, ARSH4, HIS3) were amplified from pAGE1.0^7^. The tetM gene and pUC19 *E. coli* origin were amplified from pAL1. The last 517 base pairs of the toxic gene W85.1 were amplified from pAL-Tox-2244 (Karas Lab), which is an intermediate plasmid constructed to introduce the Y676H mutation into the toxic gene.

For pNZ18-T, PCR amplicons putatively containing the ACL_0241 promoter and terminator (amplified from *A. laidlawii* PG-8A genomic DNA) were joined to the tetM CDS from pAL1 using overlap extension PCR. Specifically, each fragment was first individually PCR amplified, and products were combined into a fresh PCR reaction in a roughly 1:1:1 molar ratio, comprising 50% of the new reaction volume. The reaction was run for 15 cycles, after which 1 microliter was used as template to amplify the final stitched fragment. The resulting stitched product was cloned into pNZ18-CAH at the same location where lacZ was cloned in pNZ18-Z. To create pNZ18*-T, the mutated repB gene from pNZ18*-Z was amplified and used to replace the wild type sequence in pNZ18-T. The construct was assembled in *E. coli* strain MC1061 by co-transforming an equimolar amount of each fragment to chemically competent cells.

**PCR**

For simplex diagnostic PCRs and plasmid construction, reactions were performed using GXL PrimeSTAR (Takara Bio), with parameters according to the manufacturer’s instructions. One microliter of isolated *A. laidlawii* DNA and/or approximately 0.1 – 10 ng of plasmid DNA were used as template. For multiplex PCR, reactions were performed using Superplex polymerase (Takara Bio), with cycle parameters according to the manufacturer’s recommended multiplex protocol. One microliter of isolated DNA from yeast or *A. laidlawii* was used as template. Simplex screening primers are listed in Supplementary Table 3, and multiplex screening primers are listed in Supplementary Table 4.
